# Production of benzylglucosinolate by engineering and optimizing the biosynthetic pathway in *Saccharomyces cerevisiae*

**DOI:** 10.1101/2020.07.08.193391

**Authors:** Cuiwei Wang, Christoph Crocoll, Christina Spuur Nødvig, Uffe Hasbro Mortensen, Sidsel Ettrup Clemmensen, Barbara Ann Halkier

## Abstract

Glucosinolates are amino acid-derived defense compounds characteristic of the Brassicales order. Benzylglucosinolate (BGLS) derived from phenylalanine is associated with health-promoting effects, which has primed a desire to produce BGLS in microorganisms for a stable and rich source. In this study, we engineered the BGLS production in *Saccharomyces cerevisiae* by either stably integrating the biosynthetic genes into the genome or introducing them from plasmids. A comparison of the two approaches exhibited a significantly higher level of BGLS production (9.3-fold) by expression of the genes from genome than from plasmids. Towards optimization of BGLS production from genes stably integrated into the genome, we enhanced expression of the entry point enzymes CYP79A2 and CYP83B1 resulting in a 2-fold increase in BGLS production, but also a 4.8-fold increase in the biosynthesis of the last intermediate desulfo-benzylglucosinolate (dsBGLS). To alleviate the metabolic bottleneck in the last step converting dsBGLS to BGLS by 3’-phosphoadenosine-5’-phosphosulfate (PAPS)-dependent sulfotransferase, SOT16, we first obtained an increased BGLS production by 1.7-fold when overexpressing *SOT16*. Next, we introduced APS kinase APK1 of *Arabidopsis thaliana* for efficient PAPS regeneration, which improved the level of BGLS production by 1.7-fold. Our work shows an optimized production of BGLS in *S. cerevisiae* and the effect of different approaches for engineering the biosynthetic pathway (plasmid expression and genome integration) on the production level of BGLS.

## 1. Introduction

Glucosinolates (GLSs) are amino acid-derived defense compounds characteristic of the Brassicales order, including vegetables like broccoli, cabbages and the model plant *Arabidopsis thaliana* (Halkier and Gershenzon, 2006). Intake of brassicaceous vegetables has been linked to reduced risk of developing cancer and cardiovascular disease due to bioactivities of specific GLSs (or rather their hydrolysis products) (Traka, 2016). This has primed a desire to engineer the production of health-promoting GLSs to enable a stable and rich source as dietary supplements for the benefit of human health. Biotechnological approaches have been used to engineer and optimize the production of the desirable GLSs (Petersen et al., 2019, 2018; Wang et al., 2020).

The phenylalanine-derived benzylglucosinolate (BGLS) is one of the health-beneficial GLSs due to the hydrolysis product benzylisothiocyanate (Mitsiogianni et al., 2019). In *A. thaliana*, the BGLS biosynthetic pathway consists of seven enzymatic steps, including CYP79A2, CYP83B1, GSTF9, GGP1, SUR1, UGT74B1 and SOT16 (**Fig. 1**) (Sønderby et al., 2010). The first production of BGLS in a heterologous host was shown using transient expression in *Nicotiana benthamiana* (Geu-Flores et al., 2009). Recently, the BGLS production was engineered and optimized upon plasmid expression in *Escherichia coli* (Petersen et al., 2019). In this study, *Saccharomyces cerevisiae* was employed to engineer the BGLS biosynthetic pathway due to its GRAS status and high tolerance against harsh industrial conditions as well as advanced molecular tools for genetic modifications.

**Fig. 1.**
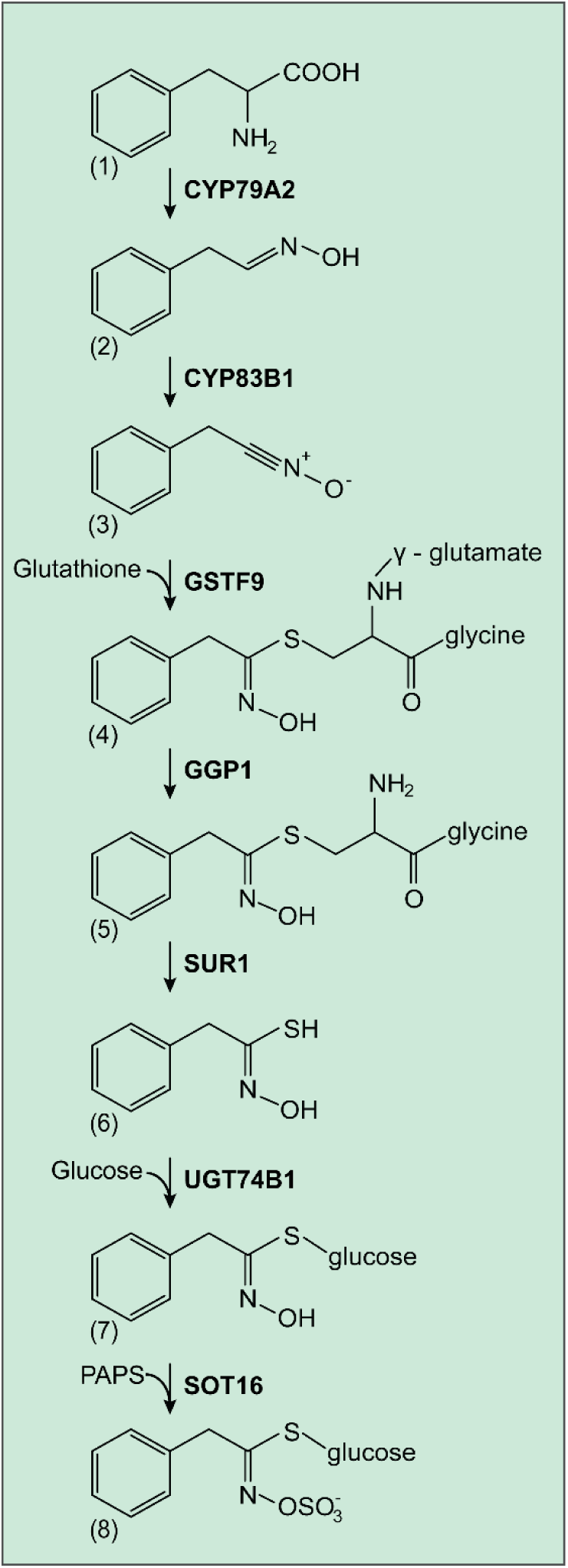
The BGLS biosynthetic pathway in *Arabidopsis thaliana*. The enzymes CYP79A2, CYP83B1, GSTF9, GGP1, SUR1, UGT74B1 and SOT16 are cytochrome P450 of the CYP79 family, cytochrome P450 of the CYP83 family, glutathione-S-transferase F9, γ-glutamyl peptidase 1, C-S lyase, UDP-glucosyltransferase 74B1 and sulfotransferase, respectively. The compounds in the BGLS pathway are 1) phenylalanine, 2) phenylacetaldoxime, 3) phenylacetonitrile oxide, 4) S-[(Z)-phenylacetohydroximoyl]-L-glutathione, 5) Cys-Gly conjugate, 6) phenylacetothiohydroximic acid, 7) desulfo-benzylglucosinolate (dsBGLS) and 8) benzylglucosinolate (BGLS).

Different approaches have been applied to express multi-gene pathways in *S. cerevisiae* either stably in genome or on plasmids. Genome integration of genes is developed upon homologous recombination. Previously, we identified 14 locations in the yeast genome for gene integration with high gene expression and without any significant impact on the growth rate (Mikkelsen et al., 2012). Each site is situated centrally in large intergenic regions of minimum 750 bp, to reduce the chance that integration influences the fitness of the strain by adversely affecting neighboring genes. Using this platform, Mikkelsen et al. (2012) demonstrated the production of tryptophan-derived indol-3-ylmethylglucosinolate in *S. cerevisiae* by stably integrating the biosynthetic genes into the genome. The platform was further advanced to a gene amplification system, CASCADE (Strucko et al., 2017), that allows amplicons containing one or more genes (i.e. entire pathways) to be efficiently introduced into the genome with defined numbers of amplicons ranging from one to nine integrated copies. For plasmid-based expression in *S. cerevisiae*, the multi-copy expression is frequently achieved via use of self-replicating, 2 μ-based plasmids (Siewers et al., 2010). The plasmids are designed to be either low, medium or high copy number plasmids (Lian et al., 2016). The 2μ-derived high-copy plasmids have been used to engineer pathways for high level expression of biosynthetic genes (Chen et al., 2013; Cheng et al., 2019; Dejong et al., 2006). Alternatively, biosynthetic pathways are commonly engineered and optimized using a combination of genome integration and plasmids (Brown et al., 2015; Galanie et al., 2015; Trenchard and Smolke, 2015).

In this study, we compared the production level of BGLS in *S. cerevisiae* upon engineering the eight-gene biosynthetic pathway by two approaches: multiple-copy plasmid expression and single-copy stable genome integration. The genome integration approach resulted in the highest yield of BGLS that was subsequently optimized by enhancing the expression level of the entry point enzymes CYP79A2 and CYP83B1 in the BGLS biosynthetic pathway. Additionally, the metabolic bottleneck in the sulfotransferase step was partially alleviated by increasing the activity of the responsible sulfotransferase enzyme and by introducing APK1 for PAPS generation.

## 2. Materials and methods

### 2.1 Generation of constructs

Four plasmids for insertion of the expression cassettes into the selected integration sites were generated according to the published method (Hansen et al., 2011). Each plasmid contains 500 – 600 bp regions of upstream and downstream of the selected integration sites on yeast chromosome XII (Mikkelsen et al., 2012). Each plasmid contains the selectable or counter selectable *Kluyveromyces lactis*-derived *URA3* marker gene that is flanked by directly repeated sequences to be recycled. The gene sequences of the BGLS biosynthetic pathway (*CYP79A2, CYP83B1, GSTF9, SUR1, GGP1, UGT74B1* and *SOT16*) and the cytochrome P450 electron-donating support gene *ATR1* were found in TAIR (The Arabidopsis Information Resource) database (https://www.arabidopsis.org/) and amplified by PCR and inserted into the plasmids for integration in pairs. A bidirectional constitutive *TEF1/PGK1*-promoter and terminators ADH1 and CYC1 were cloned into the plasmids to control the gene expression. Finally, four integration constructs were generated: pXII-1 containing *CYP79A2* and *CYP83B1*, pXII-2 containing *GSTF9* and *ATR1*, pXII-3 containing *SUR1* and *GGP1* and pXII-4 containing *SOT16* and *UGT74B1*. The detailed DNA layout was seen in **Fig. 2A**. Based on the four integration constructs, the eight genes flanked with the same promoters and terminators were cloned into the 2μ-based high-copy plasmids pESC-URA-USER and pESC-HIS (#217451, Agilent Technologies), resulting in pESC-URA-USER harboring *CYP79A2, CYP83B1, GSTF9* and *ATR1* and pESC-HIS harboring *SUR1, GGP1, SOT16* and *UGT74B1*, seen **Fig. 2B**. The plasmid pESC-URA-USER was generated in our lab by adding USER cassette based on the plasmid pESC-URA (#217454, Agilent Technologies) for USER cloning (Nour-Eldin et al., 2006).

**Fig. 2.**
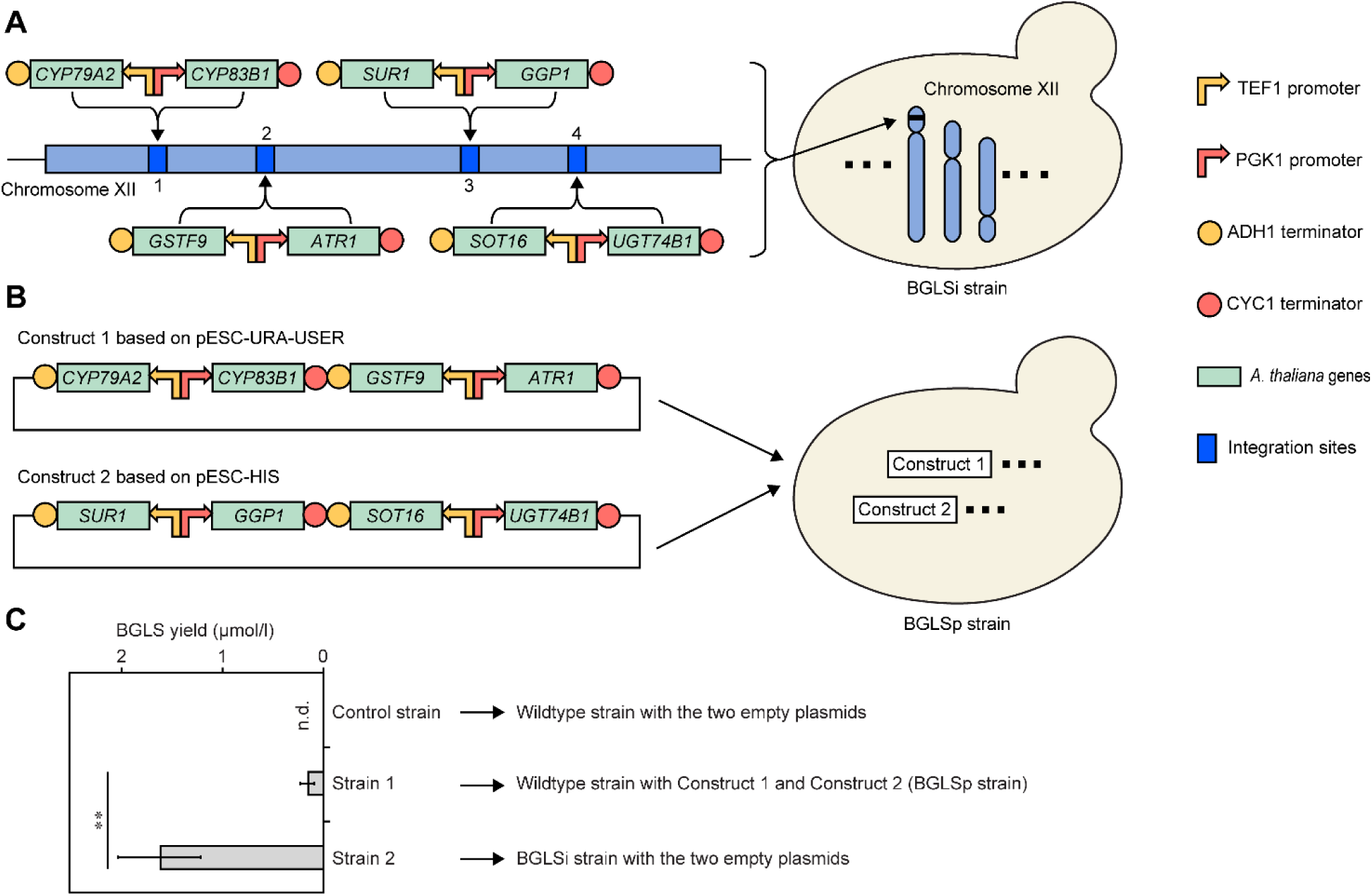
BGLS production by the *Saccharomyces cerevisiae* strains with the BGLS biosynthetic genes expressed on plasmids or by stable integration in the genome. **A)** Scheme of integration of the BGLS pathway genes in the XII chromosome of *S. cerevisiae*. The genes flanked with TEF1 promoter and ADH1 terminator or PGK1 promoter and CYC1 terminator were inserted into the integration sites in pairs. **B)** Scheme of construction of the BGLS pathway genes on plasmids. The genes are flanked with the same promoters and terminators shown in **A**. *CYP79A2, CYP83B1, GSTF9* and *ATR1* are constructed on high copy-number plasmid pESC-URA-USER and the remaining four genes are constructed on high copy-number plasmid pESC-HIS. The support gene *ATR1* is co-expressed with the BGLS pathway genes shown in **A** and **B. C)** BGLS yields. Control strain and Strain 2 respectively represent the wildtype strain and BGLSi strain transformed with the two empty plasmids pESC-URA-USER and pESC-HIS. Strain 1 (BGLSp) represents the wildtype strain transformed with the two constructs shown in **B**. The engineered strains were grown in Sc-URA-HIS-GLU media and samples were taken at 48 h. Data represent the average and standard deviation of four biological replicates. n.d. represents not detected. Exact values and *p*-value (two-sided Student’s *t*-test) are listed in **Supplementary Table S3**.

To optimize the BGLS production level, the expression cassette containing the genes *CYP79A2* and *CYP83B1* were amplified by PCR using the integration construct pXII-1 as template and constructed into the plasmid pESC-URA-USER (**Fig. 3A**). The gene *SOT16* was constructed into the plasmid pESC-HIS, flanked with galactose inducible promoter GAL1 and terminator CYC1 (**Fig. 4A**). The chloroplast localization sequences (Ravilious et al., 2012) were removed from the *APK1* gene of *A. thaliana* and the truncated gene was amplified from cDNA. The gene was constructed into the plasmid pESC-URA-USER (**Fig. 4A**). All the genes used in this study are listed in **Table 1**.

**Table 1.** The genes used to engineer and optimize BGLS production.

| Gene | Enzyme | Species | Locus |
| --- | --- | --- | --- |
| *CYP79A2* | Cytochrome P450 | *Arabidopsis thaliana* | AT5G05260 |
| *CYP83B1* | Cytochrome P450 | *Arabidopsis thaliana* | AT4G31500 |
| *GSTF9* | Glutathione S-transferase | *Arabidopsis thaliana* | AT2G30860 |
| *GGP1* | γ-glutamyl peptidase | *Arabidopsis thaliana* | AT4G30530 |
| *SUR1* | C-S lyase | *Arabidopsis thaliana* | AT2G20610 |
| *UGT74B1* | Glycosyltransferase | *Arabidopsis thaliana* | AT1G24100 |
| *SOT16* | Sulfotransferase | *Arabidopsis thaliana* | AT1G74100 |
| *ATR1* | P450 reductase | *Arabidopsis thaliana* | AT4G24520 |
| *APK1* | APS kinase | *Arabidopsis thaliana* | AT2G14750 |

**Fig. 3.**
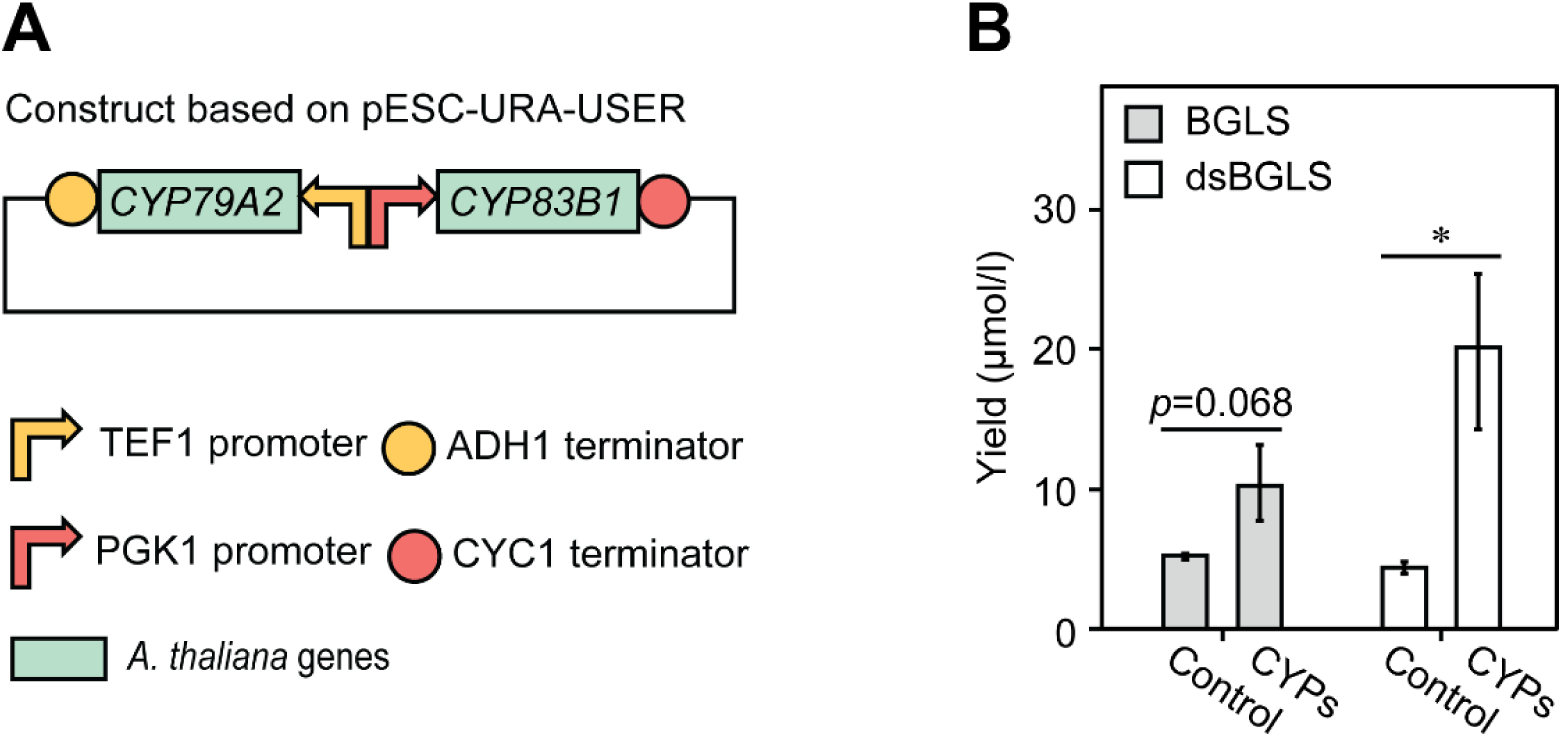
Production of BGLS and dsBGLS from the BGLSi strain by enhancing the entry point enzyme activity of CYP79A2 and CYP83B1 in the BGLS biosynthetic pathway. **A)** Construct for overexpression of *CYP79A2* and *CYP83B1. CYP79A2* and *CYP83B1* flanked with the same promoters and terminators shown in **Fig. 2B** are constructed on the high copy-number plasmid pESC-URA-USER. **B)** Yields of BGLS and dsBGLS by the BGLSi strain overexpressing the entry point genes *CYP79A2* and *CYP83B1*. Control and CYPs represent the BGLSi strain respectively containing the empty plasmid pESC-URA-USER and the construct with *CYP79A2* and *CYP83B1* shown in **A**. The engineered strains were grown in Sc-URA-GAL medium and samples were taken at 48 h. Data represent the average and standard deviation of four biological replicates. Exact values and *p*-value (two-sided Student’s *t*-test) are listed in **Supplementary Table S3**.

**Fig. 4.**
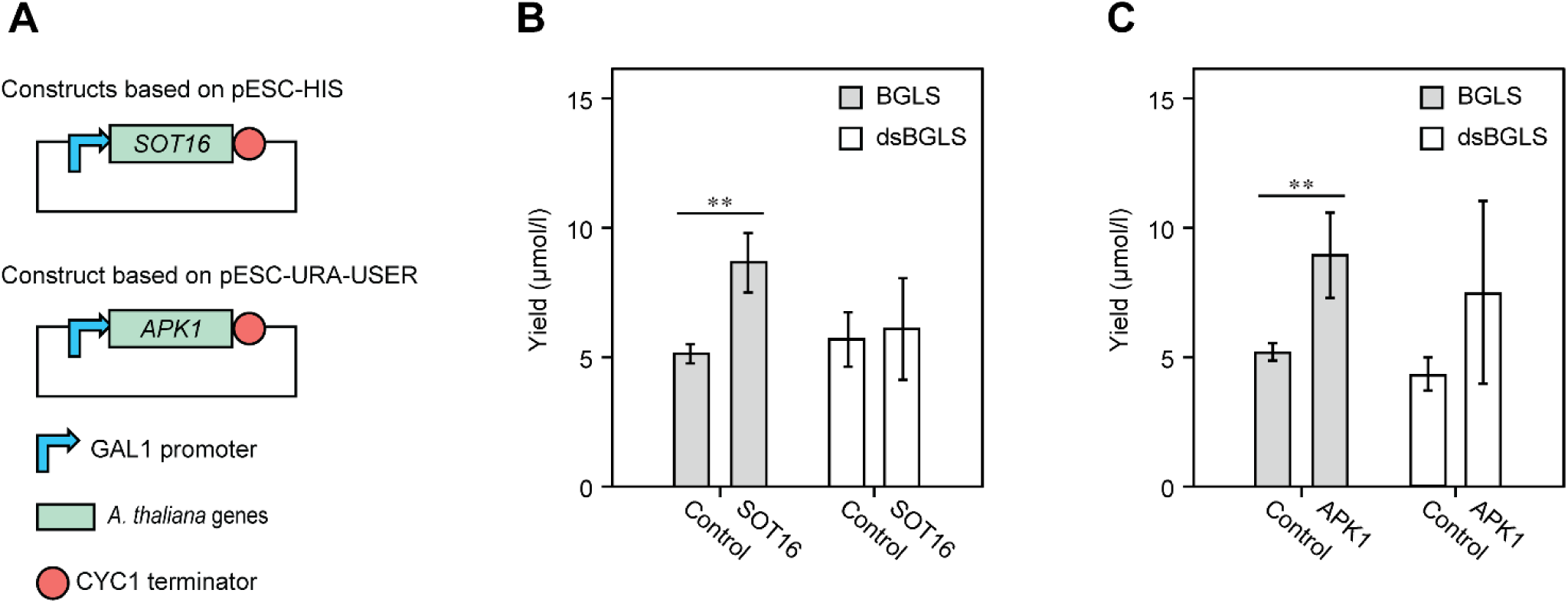
Production of BGLS and dsBGLS from the BGLSi strain by improving the last step in the BGLS biosynthetic pathway. **A)** Constructs for overexpression of *SOT16* and expression of *APK1* encoding *A. thaliana* APS kinase. Expression of all the genes are controlled by the galactose inducible promoter GAL1 and the terminator CYC1. **B)** Yields of BGLS and dsBGLS from the BGLSi strain by enhancing the enzyme activity of SOT16. The abbreviations Control and SOT16 represent the BGLSi strain respectively containing the empty plasmid pESC-HIS and the construct with *SOT16*. The engineered strains were grown in Sc-HIS-GAL medium and samples were taken at 48 h. **C)** Yields of BGLS and dsBGLS from the BGLSi strain upon expression of *APK1* for PAPS regeneration. Control and APK1 represent the BGLSi strain respectively containing the empty plasmid pESC-URA-USER and the construct with *APK1*. The engineered strains were grown in Sc-URA-GAL media and samples were taken at 48 h. Data represent the average and standard deviation of four biological replicates. In **B** and **C**, statistical significance is shown based on two-sided Student’s *t*-test. Exact values and *p*-values are listed in **Supplementary Table S3**.

### 2.2 Generation of strains

All yeast strains were generated using the previously described transformation method (Gietz and Schiestl, 2007). Transformation was confirmed by colony PCR.

To generate the yeast strain harboring the seven genes of BGLS biosynthetic pathway and the supporting gene *ATR1* on genome, the constructs pXII-1, pXII-2, pXII-3 and pXII-4 were digested by *XbaI*, respectively. The four linear DNA fragments were iteratively transformed into the yeast strain CEN.PK 113-11C. The *URA3* maker gene is eliminated by direct repeat recombination and 5-fluoroorotic acid (5-FOA) selection for the subsequent transformation (Siewers et al., 2010). The two plasmids with the eight genes for BGLS production were simultaneously transformed into CEN.PK 113-11C.

### 2.3 Growth conditions of yeast strains

All the transformants were grown on the corresponding yeast synthetic complete drop-out media with 20 g/l agar and 20g/l glucose as carbon source according to their harboring plasmids. For generation of the yeast strain containing the BGLS biosynthetic genes on genome, the transformants were grown on synthetic complete uracil drop-out (Sc-URA) media with 1.92 g/l Yeast Synthetic Drop-out Medium Supplements without uracil (Sigma-Aldrich) and 6.7 g/l Yeast Nitrogen Base Without Amino Acids (Sigma-Aldrich). Subsequently, the strains were grown on synthetic complete media containing 30 mg/l uracil and 740 mg/l 5-FOA to excise the *URA3* maker gene. For transformation of pESC-URA-USER-based constructs and pESC-HIS-based constructs, the transformants were grown on Sc-URA media and synthetic complete histidine drop-out (Sc-HIS) media, respectively. Similarly, the synthetic complete uracil and histidine drop-out (Sc-URA-HIS) media were used to select the yeast strain transformed with both pESC-URA-USER-based and pESC-HIS-based constructs.

All the strains were cultured in 100 ml baffled flasks containing 30 ml media at 30 °C, 150 rpm. Glucose was used as carbon source to grow all the pre-cultures and the first BGLS-producing cultures for comparison of genome integration and plasmid-based introduction of the BGLS biosynthetic genes. Galactose was used as carbon source to grow the BGLS-producing culture for optimization of BGLS production. Single colonies were inoculated in media for around 24 h as pre-culture. The proper volume of preculture was precipitated at 5,000 × *g* for 5 min to collect the cells, resulting in the original OD_600_ to 0.6 in fresh media as expression culture.

### 2.4 Metabolite extraction and LC-MS analysis

The expression culture was precipitated at 17,000 × *g* for 5 min to collect the supernatant. The supernatant was diluted by 10-fold with a buffer containing 10 μg/ml 13C-, 15N-labelled amino acids (Algal amino acids 13C, 15N, Isotec, Miamisburg, US) and 2 μmol/l sinigrin (PhytoLab, Vestenbergsgreuth, Germany). Subsequently, the diluted samples were filtered (Durapore® 0.22 μm PVDF filters, Merck Millipore, Tullagreen, Ireland) and used directly for LC-MS analysis. Sinigrin and 13C, 15N-Phe were used as internal standard to quantify BGLS and dsBGLS, respectively.

BGLS, dsBGLS and the level of amino acids were monitored by LC-MS analysis using a modified version of previously described method (Mirza et al., 2016). Briefly, an Advance UHPLC system (Bruker, Bremen, Germany) was used for chromatography. A Zorbax Eclipse XDB-C18 column (100×3.0 mm, 1.8 μm, Agilent Technologies, Germany) was used for separation. The mobile phases A and B were formic acid (0.05% (v/v)) in water and acetonitrile, respectively. The elution profile was as follows: 0-1.2 min 3% B; 1.2-4.3 min 3-65% B; 4.3-4.4 min 65-100% B; 4.4-4.9 min 100% B, 4.9-5.0 min 100-3% B and 5.0-6.0 min 3% B. Mobile phase flow rate was 500 μl/min with the column temperature maintained at 40 °C. The liquid chromatography was coupled to an EVOQ Elite TripleQuad mass spectrometer (Bruker, Bremen, Germany) equipped with an electrospray ionization source (ESI). Infusion experiments with pure standards were used to optimize instrument parameters. Ionspray voltage was set to 3000 V or −4000 V in positive and negative ionization mode, respectively. Cone temperature was maintained at 350 °C and cone gas (nitrogen) flow to 20 psi. Heated probe temperature was maintained at 400 °C and probe gas flow to 50 psi. Nebulising gas (nitrogen) was set to 60 psi and collision gas (argon) to 1.6 mTorr.

Multiple reaction monitoring (MRM) was used to monitor analyte parent ion to product ion transitions: MRM transitions for 13C-, 15N-labelled phenylalanine was used as previously reported (Docimo et al., 2012). MRM transitions for BGLS, dsBGLS and sinigrin were used as previously reported (Crocoll et al., 2016). Q1 and Q3 quadrupoles were maintained at unit resolution. Data acquisition and processing were achieved by Bruker MS Workstation software (Version 8.2.1, Bruker, Bremen, Germany). For analytes, multiple transitions were monitored and the transition used for quantification is marked as quantifier (Qt). The information of transitions and collision energies is seen in **Supplementary Table S1**. Dilution series of the respective analytes were used to calculate response factors to the respective internal standards. The selection of internal standards is based on matching ionization mode with the analyte of interest (i.e. BGLS in negative ionization mode; dsBGLS and amino acids in positive ionization mode). Due to matrix effect from the cultures on the quantification of the analytes, the correction factors were calculated into the response factors in **Supplementary Table S2**.

## 3. Results

### 3.1 Engineering BGLS production in S. cerevisiae

The seven-gene BGLS biosynthetic pathway – *CYP79A2, CYP83B1, GSTF9, GGP1, SUR1, UGT74B1* and *SOT16* – was engineered in *S. cerevisiae* CEN.PK 113-11C. The gene *ATR1* encoding NADPH cytochrome P450 reductase from *A. thaliana* was co-expressed to support the function of cytochrome P450 enzymes. The eight genes driven by the strong constitutive promoters TEF1 or PGK1 were pairwise stably integrated into four well-characterized sites on the chromosome XII (Mikkelsen et al., 2012) and this strain was named BGLSi strain (**Fig. 2A**). In parallel, we generated two 2μ-derived high-copy constructs containing the BGLS biosynthetic genes under the control of the same promoters and terminators with the ones for construction of the BGLSi strain (**Fig. 2B**). Subsequently, the two constructs were transformed into CEN.PK 113-11C, resulting in a yeast strain named BGLSp strain (**Fig. 2B**).

To compare the effect of gene expression way (single-copy genome integration or high-copy plasmids) on BGLS production, we additionally transformed the BGLSi strain with the two empty plasmids that were used for expression of the BGLS pathway genes in the BGLSp strain. When the two strains (the BGLSi strain with the two empty plasmids and the BGLSp strain) were grown in glucose-containing synthetic media minus uracil and histidine for 48 h, we found that both strains produced BGLS (**Fig. 2C**), which shows that the heterologous expression of the BGLS biosynthetic genes in both ways was functional in *S. cerevisiae*. The BGLSi strain with the two empty plasmids produced 1.67 µmol/l BGLS, and the BGLSp produced 0.18 µmol/l (**Fig. 2C** and **Supplementary Table S3**). This means that expression of the BGLS biosynthetic genes through stable genome integration results in 9.3-fold higher production level of BGLS compared to expression from the plasmids despite that the copy number of plasmids is higher than genome integration.

### 3.2 Optimization of BGLS production by enhancing the entry point of the BGLS biosynthetic pathway

The low production level of BGLS from the BGLSi strain primed us to optimize the production for higher yield. First, we boosted the expression of the entry point enzymes through introduction of *CYP79A2* and *CYP83B1* driven by the constitutive promoters TEF1 and PGK1 from a high-copy plasmid (**Fig. 3A**). We grew the strains in galactose-containing synthetic media minus uracil for 48 h and found that similar levels of BGLS (5.21 µmol/l) and dsBGLS (4.43 µmol/l) were produced by the control strain, the BGLSi strain with the empty plasmid pESC-URA-USER (**Fig. 3B** and **Supplementary Table S3**). The amount of BGLS increased by 2-fold with the level at 10.37 µmol/l while a 4.8-fold increase in the production of dsBGLS (20.67 µmol/l) was observed through boosting expression of the entry point enzymes (**Fig. 3B** and **Supplementary Table. S3**). This result shows that enhanced expression level of the entry point enzymes CYP79A2 and CYP83B1 increases the BGLS production level and the flux through the pathway. However, the large accumulation of dsBGLS indicates that the last step of the BGLS pathway is a metabolic bottleneck under this condition.

### 3.3 Alleviation of the metabolic bottleneck at the sulfotransferase last step in the BGLS biosynthetic pathway

Towards alleviating the metabolic bottleneck at the last step in the BGLS pathway, we first enhanced the expression of the responsible gene *SOT16* by increasing copy number and using the strong galactose inducible promoter GAL1 to regulate the expression (**Fig. 4A**). We monitored the production level of BGLS and dsBGLS at 48 h after induction and found that the BGLS yield significantly increased by 1.7-fold to 9.12 µmol/l when expression of *SOT16* was enhanced (**Fig. 4B**). However, no significant difference in dsBGLS accumulation was observed by overexpression of *SOT16* (**Fig. 4B**), suggesting that elevated expression level of *SOT16* improves BGLS production but cannot overcome the metabolic bottleneck. The results suggest that availability of the PAPS co-factor for the sulfotransferase may be a limiting factor.

To increase PAPS availability, we introduced a truncated version of *A. thaliana* APS kinase APK1 without chloroplast signal peptides to boost PAPS generation. We introduced the gene driven by the galactose-inducible promoter GAL1 on the high-copy plasmid pESC-URA-USER (**Fig. 4A**). Expression of the truncated *APK1* gene resulted in an increased production of BGLS by 1.7-fold but the accumulation level of dsBGLS did not significantly change (**Fig. 4C**). The result demonstrates that the last sulfotransferase step is still a metabolic bottleneck even though introduction of APK1 increases the BGLS production.

## 4. Discussion

In this study, we used *S. cerevisiae* as cell factory to engineer BGLS production using either genome integration or plasmid-based expression of the biosynthetic genes. Subsequently, genome integration-based BGLS production level was improved through increased expression of the entry point enzymes (CYP79A2 and CYP83B1). Additionally, a metabolic bottleneck at the last enzymatic step catalyzed by the sulfotransferase SOT16 was partly alleviated by overexpression of *SOT16* and introduction of APK1 for the regeneration of co-factor PAPS.

The eight BGLS biosynthetic genes were introduced into *S. cerevisiae* by two engineering patterns, single-copy genome integration and multi-copy plasmid expression. We expected to obtain higher production level of BGLS when introducing the pathway on high-copy plasmids, but the single-copy genome integration resulted in almost 10-fold higher production (**Fig. 2**). A possible explanation for this is low genetic stability of plasmids in yeast. For example, two fluorescent proteins, CFP and RFP, expressed from 2μ-based plasmids showed that only 54% of the cells produced both proteins at mid-exponential phase (Strucko et al., 2017). Another study reported that only around 28% of the yeast cells transformed with two 2μ-based plasmids containing three genes for synthesizing artemisinic acid maintained both of the plasmids after 120 h growth (Ro et al., 2008). Furthermore, copy numbers affect plasmid stability, evidenced by an assay upon expressing cheilanthifoline synthase from *Eschscholzia californica*: 90% of the low-copy plasmid was retained while only 10.7% of the high-copy plasmid was maintained (Trenchard and Smolke, 2015). However, increasing copy number of expressed genes is usually used to increase the enzyme level and thus improve the production level of target products (Chen et al., 2012; Ro et al., 2008). Therefore, for the future work multi-copy integration of the pathway into the genome would be a promising way for higher production level of BGLS.

Expectedly, the metabolic bottleneck at the last step as evidenced by the accumulation of dsBGLS in media was enhanced when flux through the pathway was increased by overexpression of the entry point genes, *CYP79A2* and *CYP83B1* (**Fig. 3**). Possible explanations of the bottleneck could be inefficient enzyme level of sulfotransferase SOT16 or inadequate supply of co-factor PAPS. However, the bottleneck still existed when the expression level of *SOT16* and PAPS generation were increased even though the strategies enabled higher amount of BGLS (**Fig. 4B** and **C**). These approaches did not completely overcome the problem of the dsBGLS accumulation in the last step, suggesting that PAPS supply for conversion of dsBGLS into BGLS is still a metabolic bottleneck. It has been reported that the intracellular levels of PAPS is fine-tuned in the microorganisms due to toxicity (Russel et al., 1990). Moreover, when PAPS has donated a sulfate group to dsBGLS, adenosine-3’,5’-bisphosphate (PAP) accumulates, and accumulation of PAP has been reported to inhibit sulfotransferases and be toxic to yeast (Burkart et al., 2000). Alternatively, export of dsBGLS from the cells could be a means of detoxification leading to premature abortion of the BGLS pathway.

Noticeably, the problem of dsBGLS accumulation was also observed when engineering BGLS production in *E. coli* and *N. benthamiana* (Petersen et al., 2019; Møldrup et al., 2011). Towards alleviating the problem, the sulfur metabolism was modified in each BGLS-producing host. Co-expression of APS kinase APK2 reduced the majority of dsBGLS accumulation and increased the BGLS level 16-fold in *N. benthamiana* (Møldrup et al., 2011). However, in *E. coli* overexpression of three native PAPS-generating genes encoding for an adenylyl-sulfate kinase and two subunits of a sulfate adenylyltransferase did not increase and even reduced the BGLS production level with supplemented different sulfur sources in media (Petersen et al., 2019). Alternative sulfotransferase enzymes for the last step of the BGLS pathway were tested in the two organisms but the BGLS production level was not improved (Møldrup et al., 2011; Petersen et al., 2019). Additionally, the increased expression of *SOT16* did not assist the dsBGLS to BGLS conversion in *E. coli* (Petersen et al., 2019), whereas overexpression of SOT16 increasing the BGLS yield in yeast from our study. The BGLS production have been previously engineered in *E. coli* with the highest level being 20.3 μmol/L (equivalent to 8.3 mg/L) (Petersen et al., 2019). The highest yield of BGLS produced by the engineered yeast strain is 10.37 µmol/L (= 4.2 mg/L). This could be caused by many differences between the eukaryotic *S. cerevisiae* and prokaryotic *E. coli* – one of which is that the endogenous cytochrome P450s of *S. cerevisiae* could be involved in the metabolism of the intermediates in the BGLS biosynthetic pathway (Crešnar and Petrič, 2011) whereas *E. coli* does not have this problem due to the absence of the native cytochrome P450s.

In summary, this study reports the first comparison of the production of the plant natural product in yeast – BGLS – from a multi-gene pathway expressed on high-copy plasmids or by stable integration in genome. This 9.3-fold higher production shows that genome integration is the preferred strategy. Fortunately, new genome editing technologies such as e.g. Crispr-Cas9 has greatly improved the speed and accuracy of genetic engineering.

## Author contributions

CW, CC and BAH designed this study with support by UHM and wrote the manuscript based on a draft by CW. CW performed the experiments except the BGLSi strain constructed by CSN under supervision of UHM. SEC helped CW to grow some cultures. CC performed LC-MS method development and analysis.

## Acknowledgements

This work was supported by the Danish National Research Foundation [DNRF99] and the Novo Nordisk Foundation [NNF17OC0027710].

## Notes

### Competing Interest Statement

The authors have declared no competing interest.

## References

Brown, S., Clastre, M., Courdavault, V., O’Connor, S.E., 2015. De novo production of the plant-derived alkaloid strictosidine in yeast. Proc. Natl. Acad. Sci. USA 112, 3205–3210. https://doi.org/org/10.1073/pnas.1423555112.

Burkart, M.D., Izumi, M., Chapman, E., Lin, C.H., Wong, C.H., 2000. Regeneration of PAPS for the enzymatic synthesis of sulfated oligosaccharides. J. Org. Chem. 65, 5565–5574. https://doi.org/10.1021/jo000266o.

Chen, Y., Daviet, L., Schalk, M., Siewers, V., Nielsen, J., 2013. Establishing a platform cell factory through engineering of yeast acetyl-CoA metabolism. Metab. Eng. 15, 48–54. https://doi.org/10.1016/j.ymben.2012.11.002.

Chen, Y., Partow, S., Scalcinati, G., Siewers, V., Nielsen, J., 2012. Enhancing the copy number of episomal plasmids in *Saccharomyces cerevisiae* for improved protein production. FEMS Yeast Res 12, 598–607. https://doi.org/10.1111/j.1567-1364.2012.00809.x.

Cheng, S., Liu, X., Jiang, G., Wu, J., Zhang, J.-L., Lei, D., Yuan, Y.-J., Qiao, J., Zhao, G.-R., 2019. Orthogonal Engineering of Biosynthetic Pathway for Efficient Production of Limonene in *Saccharomyces cerevisiae*. ACS Synth. Biol. 8, 968–975. https://doi.org/10.1021/acssynbio.9b00135.

Crešnar, B., Petrič, S., 2011. Cytochrome P450 enzymes in the fungal kingdom. Biochim. Biophys. Acta 1814, 29–35. https://doi.org/10.1016/j.bbapap.2010.06.020.

Crocoll, C., Halkier, B.A., Burow, M., 2016. Analysis and quantification of glucosinolates. Curr. Protoc. Plant Biol. 1, 385–409. https://doi.org/10.1002/cppb.20027.

Dejong, J.M., Liu, Y., Bollon, A.P., Long, R.M., Jennewein, S., Williams, D., Croteau, R.B., 2006. Genetic engineering of taxol biosynthetic genes in *Saccharomyces cerevisiae*. Biotechnol. Bioeng. 93, 212–224. https://doi.org/10.1002/bit.20694.

Docimo, T., Reichelt, M., Schneider, B., Kai, M., Kunert, G., Gershenzon, J., D’Auria, J.C., 2012. The first step in the biosynthesis of cocaine in Erythroxylum coca: the characterization of arginine and ornithine decarboxylases. Plant Mol. Biol. 78, 599–615. https://doi.org/10.1007/s11103-012-9886-1.

Galanie, S., Thodey, K., Trenchard, I.J., Filsinger Interrante, M., Smolke, C.D., 2015. Complete biosynthesis of opioids in yeast. Science. 349, 1095–1100. https://doi.org/10.1126/science.aac9373.

Geu-Flores, F., Nielsen, M.T., Nafisi, M., Møldrup, M.E., Olsen, C.E., Motawia, M.S., Halkier, B.A., 2009. Glucosinolate engineering identifies a gamma-glutamyl peptidase. Nat. Chem. Biol. 5, 575–577. https://doi.org/10.1038/nchembio.185.

Gietz, R.D., Schiestl, R.H., 2007. Frozen competent yeast cells that can be transformed with high efficiency using the LiAc/SS carrier DNA/PEG method. Nat. Protoc. 2, 1–4. https://doi.org/10.1038/nprot.2007.17.

Halkier, B.A., Gershenzon, J., 2006. Biology and biochemistry of glucosinolates. Annu. Rev. Plant Biol. 57, 303–333. https://doi.org/10.1146/annurev.arplant.57.032905.105228.

Hansen, B.G., Salomonsen, B., Nielsen, M.T., Nielsen, J.B., Hansen, N.B., Nielsen, K.F., Regueira, T.B., Nielsen, J., Patil, K.R., Mortensen, U.H., 2011. Versatile enzyme expression and characterization system for Aspergillus nidulans, with the Penicillium brevicompactum polyketide synthase gene from the mycophenolic acid gene cluster as a test case. Appl. Environ. Microbiol. 77, 3044–3051. https://doi.org/10.1128/AEM.01768-10.

Lian, J., Jin, R., Zhao, H., 2016. Construction of plasmids with tunable copy numbers in Saccharomyces cerevisiae and their applications in pathway optimization and multiplex genome integration. Biotechnol. Bioeng. 113, 2462–2473. https://doi.org/10.1002/bit.26004.

Mikkelsen, M.D., Buron, L.D., Salomonsen, B., Olsen, C.E., Hansen, B.G., Mortensen, U.H., Halkier, B.A., 2012. Microbial production of indolylglucosinolate through engineering of a multi-gene pathway in a versatile yeast expression platform. Metab. Eng. 14, 104–111. https://doi.org/10.1016/j.ymben.2012.01.006.

Mirza, N., Crocoll, C., Erik Olsen, C., Ann Halkier, B., 2016. Engineering of methionine chain elongation part of glucoraphanin pathway in *E. coli*. Metab. Eng. 35, 31–37. https://doi.org/10.1016/j.ymben.2015.09.012.

Mitsiogianni, M., Koutsidis, G., Mavroudis, N., Trafalis, D.T., Botaitis, S., Franco, R., Zoumpourlis, V., Amery, T., Galanis, A., Pappa, A., Panayiotidis, M.I., 2019. The Role of Isothiocyanates as Cancer Chemo-Preventive, Chemo-Therapeutic and Anti-Melanoma Agents. Antioxidants (Basel) 8. https://doi.org/10.3390/antiox8040106.

Møldrup, M.E., Geu-Flores, F., Olsen, C.E., Halkier, B.A., 2011. Modulation of sulfur metabolism enables efficient glucosinolate engineering. BMC Biotechnol. 11, 12. https://doi.org/10.1186/1472-6750-11-12.

Nour-Eldin, H.H., Hansen, B.G., Nørholm, M.H.H., Jensen, J.K., Halkier, B.A., 2006. Advancing uracil-excision based cloning towards an ideal technique for cloning PCR fragments. Nucleic Acids Res. 34, e122. https://doi.org/org/10.1093/nar/gkl635.

Petersen, A., Crocoll, C., Halkier, B.A., 2019. De novo production of benzyl glucosinolate in Escherichia coli. Metab. Eng. 54, 24–34. https://doi.org/10.1016/j.ymben.2019.02.004.

Petersen, A., Wang, C., Crocoll, C., Halkier, B.A., 2018. Biotechnological approaches in glucosinolate production. J. Integr. Plant Biol. 60, 1231–1248. https://doi.org/10.1111/jipb.12705.

Ravilious, G.E., Nguyen, A., Francois, J.A., Jez, J.M., 2012. Structural basis and evolution of redox regulation in plant adenosine-5’-phosphosulfate kinase. Proc. Natl. Acad. Sci. USA 109, 309–314. https://doi.org/10.1073/pnas.1115772108.

Ro, D.-K., Ouellet, M., Paradise, E.M., Burd, H., Eng, D., Paddon, C.J., Newman, J.D., Keasling, J.D., 2008. Induction of multiple pleiotropic drug resistance genes in yeast engineered to produce an increased level of anti-malarial drug precursor, artemisinic acid. BMC Biotechnol. 8, 83. https://doi.org/10.1186/1472-6750-8-83.

Russel, M., Model, P., Holmgren, A., 1990. Thioredoxin or glutaredoxin in *Escherichia coli* is essential for sulfate reduction but not for deoxyribonucleotide synthesis. J. Bacteriol. 172, 1923–1929. https://doi.org/10.1128/jb.172.4.1923-1929.1990.

Siewers, V., Mortensen, U., Nielsen, J., 2010. Genetic engineering tools for *Saccharomyces cerevisiae*. In: Baltz, R., Davies, J.E., Demain, A.L. (Eds.), Manual of Industrial Microbiology and Biotechnology. ASM Press, Washington, DC, USA, pp.287–301. https://doi.org/10.1128/9781555816827.ch20.

Sønderby, I.E., Geu-Flores, F., Halkier, B.A., 2010. Biosynthesis of glucosinolates--gene discovery and beyond. Trends Plant Sci. 15, 283–290. https://doi.org/10.1016/j.tplants.2010.02.005.

Strucko, T., Buron, L.D., Jarczynska, Z.D., Nødvig, C.S., Mølgaard, L., Halkier, B.A., Mortensen, U.H., 2017. CASCADE, a platform for controlled gene amplification for high, tunable and selection-free gene expression in yeast. Sci. Rep. 7, 41431. https://doi.org/10.1038/srep41431.

Traka, M.H., 2016. Health benefits of glucosinolates, in: Glucosinolates, Advances in Botanical Research. Elsevier, pp. 247–279. https://doi.org/10.1016/bs.abr.2016.06.004.

Trenchard, I.J., Smolke, C.D., 2015. Engineering strategies for the fermentative production of plant alkaloids in yeast. Metab. Eng. 30, 96–104. https://doi.org/10.1016/j.ymben.2015.05.001.

Wang, C., Crocoll, C., Agerbirk, N., Halkier, B.A., 2020. Engineering and optimization of the 2-phenylethylglucosinolate production in *Nicotiana benthamiana* by combining biosynthetic genes from *Barbarea vulgaris* and *Arabidopsis thaliana*. BioRxiv. https://doi.org/10.1101/2020.05.12.090720.

